# Cleavage, down-regulation and aggregation of serum amyloid A

**DOI:** 10.1101/811398

**Authors:** Wenhua Wang, Prabir Khatua, Ulrich H.E. Hansmann

## Abstract

Various diseases cause over-expression of the serum amyloid A protein (SAA), which leads in some, but not all, cases to amyloidosis as a secondary disease. The response to the over-expression involves dissociation of SAA hexamer and subsequent cleavage of the released monomers, most commonly yielding fragments SAA_1−76_ of the full-sized SAA_1−104_. We report results from molecular dynamic simulations that probe the role of this cleavage for down-regulating activity and the concentration of SAA. We propose a mechanism that relies on two elements. First, the probability to assemble into hexamers is lower for the fragments than it is for the full-sized protein. Second, unlike other fragments SAA_1−76_ can switch between two distinct configurations. The first kind is easy to proteolyze (allowing a fast reduction of SAA concentration) but prone to aggregation, while the situation is opposite for the second kind. If the time scale for amyloid formation is longer than the one for proteolysis, the aggregation-prone species dominates. However, if environmental conditions such as low pH increase the risk of amyloid formation, the ensemble shifts toward the more protected form. We speculate that SAA amyloidosis is a failure of the switching mechanism leading to accumulation of the aggregation-prone species and subsequent amyloid formation.

---

In order to function properly, a protein has to take a specific structure, either by itself or in complex with other molecules. Misfolded proteins lose their function and are usually degraded in cells by proteolytic cleavage.^1^ In general much longer time scales are required for a competing process by which unfolded or misfolded proteins aggregate instead into assemblies characterized by a cross-beta structure termed as “amyloid”. While these amyloids have sometimes specific functions in organisms (for instance as storage of hormones^2^ or as a matrix in bacterial biofilms^3^), their presence is in humans and other mammals more often the hallmark of neurodegenerative, metabolic and other diseases.^4–6^

These amyloid diseases are in some cases secondary illnesses. For instance, the 104-residue long serum amyloid A protein (SAA) is implied in the transport of cholesterol in high-density lipoprotein (HDL) particles, and thought to play also a role in the regulation of inflammation. In its active form, the protein assembles as a hexamer, built from two layers of trimers. The structure of the monomer has been resolved by X-ray crystallography and deposited in the Protein Data Bank (PDB-ID: 4IP9). The four-helix bundle is shown in Fig. 1(a) and consists of the N-terminal helix-I formed by residues 1–27, helix-II by residues 32–47, helix-III by residues 50–69, and the C-terminal helix-IV by residues 73–88. The structure of SAA changes little when part of the hexamer (PDB-ID 4IP8),^7^ where the chains in each trimer are packed together by the N-terminal helices, see Fig. 1(b), and cholesterol binds at the interface between the two trimers. Diseases such as colon cancer, inflammatory bowel disease, or rheumatoid arthritis can cause over-expression of SAA. The resulting serum concentration of about 1 mg/mL^8^ is more than 1000 times higher than in healthy persons and gives rise in some patients to the outbreak of SAA amyloidosis, which is characterized by the appearance of amyloid deposits, most commonly in liver, spleen, and kidney. As the deposits may interfere with the function of the affected organs, they add to the symptoms of the primary disease.^9^ A drastic example is the failure of renal function and subsequent death in captive cheetahs caused by amyloid deposits, which themselves are due to over-expression of SAA as a consequence of stress-related inflammatory diseases.^10,11^

**Figure 1:**
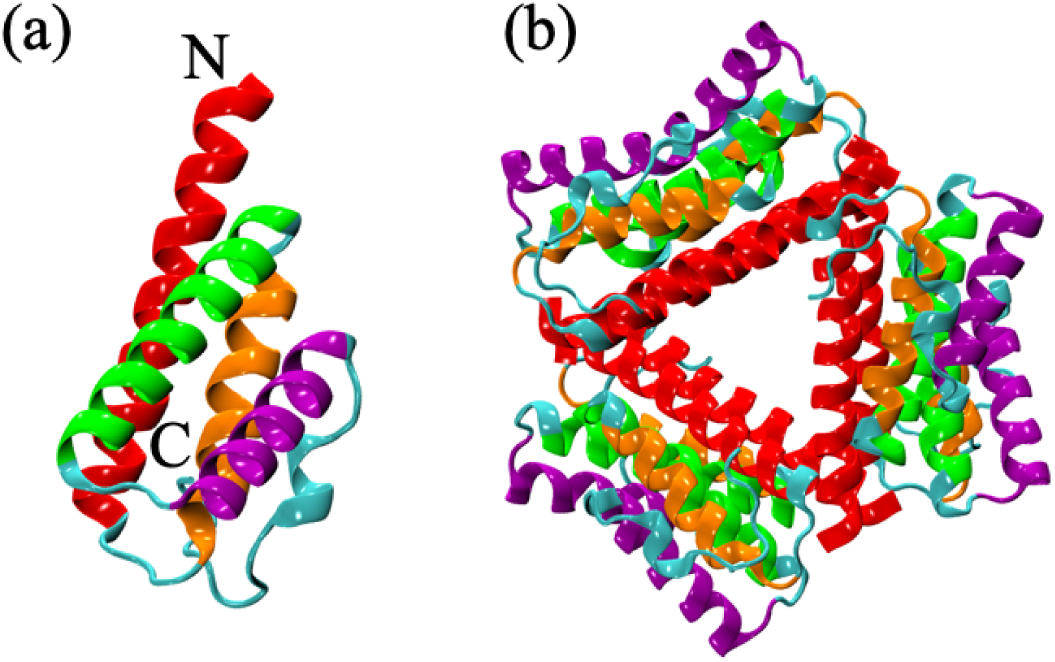
Crystal structure model of the full-sized serum amyloid A (SAA_1*−*104_) (a) monomer (PDB-ID: 4IP9) and (b) hexamer (PDB-ID:4IP8) as deposited in the Protein Data Bank. The four helices are colored as follows: helix-I (red), helix-II (orange), helix-III (green), and helix-IV (magenta).

Since SAA amyloidosis is not observed in all patients with the primary disease, it cannot be caused solely by crowding due to the high concentration. More likely it is a failure of a mechanism to down-regulate the HDL transport and other functionality after SAA over-expression, and/or to regulate the immune response to the primary disease.^12^ However, this regulation mechanism is not fully understood. Binding with glycosaminoglycans (GAGs), such as heparin/heparan sulfate (HS) leads to dissociation of the hexamer.^13,14^ This process should by itself foster down-regulation of SAA activity; however, the released 104-residue SAA_1−_104 monomers are in a second step also cleaved into smaller fragments of 45 to 94 residues.^15^ Most commonly found in amyloid deposits is the truncated fragment SAA_1−76_,^16^ but the only resolved fibril model is for the even smaller fragment SAA_2−55_ (PDB-ID: 6MST).^17^ It is known from mutation experiments that the first eleven N-terminal residues are crucial for SAA misfolding and aggregation.^18^ As part of a hexamer, in the native structure, this segment is cached in helix-I (residues 1–27) and stabilized by interactions with the neighboring chains, but in isolated monomers and fragments, it may be flexible enough to form strand-like segments.^19^ This was observed in our lab in molecular dynamics simulations of the fragment SAA_1−13_ where the segment alternated between an *α*-helix and a *β*-hairpin.^20^ Nordling et al^21^ have proposed that by taking such strand-like configurations, the N-terminal segment could nucleate fibril formation. Given the raised risk for amyloid formation (and subsequent amyloidosis), the question arises for what reason the full-length SAA proteins are cleaved into smaller fragments. In the present paper we have explored this question using molecular dynamics simulations of the full-length SAA protein and various fragments, both as monomers and assembled into a hexamer. We establish that the cleavage contributes to the down-regulation of SAA activity by shifting the equilibrium from hexamers toward monomers, thereby reducing SAA concentration.

We hypothesize that SAA_1−76_ is the most commonly found fragment because unlike smaller or larger fragments it allows switching between two distinct structures, enabling a fine-tuning of the response to SAA over-expression. Dominant at neutral pH is a structure (coined by us helix-weakened) that allows for easy degradation, helping therefore to lower quickly the SAA concentration. However, the first eleven N-terminal residues also have an increased risk in helix-weakened configurations to unfold from helix-I, and to form strand-like segments which in turn may nucleate amyloid growth. If acidic conditions raise the risk for aggregation and amyloid formation, the equilibrium shifts toward an alternative configuration (termed by us helix-broken) where the N-terminus is more stable, but which is more difficult to degrade. We speculate that in most patients where colon cancer, inflammatory bowel disease, or rheumatoid arthritis lead to over-expression of SAA, the described mechanism down-regulates activity and concentration of SAA, but that if the switching mechanism is impeded or over-whelmed, an over-abundance of the more aggregation-prone helix-weakened configurations gives rise to SAA amyloidosis.

## Materials and methods

### Initial conformations

For the full-sized serum amyloid A protein SAA_1−104_, we use in our simulations as start configurations the crystal structures, derived by X-ray crystallography and deposited in the Protein Data Bank (PDB) under identifiers 4IP8 (monomer) and 4IP9 (hexamer). Removing residues 77-104 from these two structures leads to our initial configurations for the fragment SAA_1−76_. Each of the two monomers is placed in the center of a cubic box of edge length 6.8 nm, filled with ∼ 10,000 water molecules; while for the two hexamer systems, the cubic box has an edge size of 9.8 nm and is filled with ∼ 28,000 water molecules. In a similar way, we also generate from the SAA monomer two shorter fragments SAA_1−27_ and SAA_1−47_ that are put again into cubic boxes with edge size 6.2 nm (SAA_1−27_) and 6.8 nm (SAA_1−47_), filled with ∼ 7,800 and ∼ 10,000 water molecules, respectively.

### Simulation protocol

Our molecular dynamics simulations are performed by using the GROMACS 2018.2 software package.^22^ Protein-protein and protein-water interactions are modeled with the CHARMM 36m force field^23^ and the TIP3P^24^ water model, a combination that is known to perform well in simulations of amyloid assembly.^25^ We use the LINCS algorithm^26^ to constrain the bond length and the SETTLE algorithm^27^ to maintain water geometry. As periodic boundary conditions are selected, we use the Particle Mesh Ewald (PME) algorithm^28^ to calculate the electrostatic interaction with a cut-off distance of 1.2 nm, the default value in GROMACS for electrostatic and van der Waals (vdW) interactions.

The molecular dynamics simulations are performed in an isothermal-isobaric (NPT) ensemble, with temperature set to 310 K by a v-rescale thermostat,^29^ and pressure set to 1 bar by a Parrinello-Rahman barostat.^30^ The equations of motions are integrated with a time step of 2 fs. Each system is followed over three independent trajectories of either 1 *µ*s (monomers) or 500 ns (hexamers) duration, starting from distinct initial configurations generated by introducing random noise to the respective coordinates and velocities. Measurements are taken every 50 ps and stored for further analysis.

### Observable

Time evolution of structures is followed by calculating the root-mean-square-deviation (RMSD) to the start configuration using our in-house code. Similarly, we measured the solvent accessible surface area (SASA) and the cavity diameter <*d_cavity_*>) with our in-house codes. The later quantity is approximated by averaging over the center-of-mass distances between the N-terminal helix-I segments of adjacent units of both layers. This approximation is justified as each of the two trimer layers (see figure Fig. 1(b)) is formed from the helices in the respective chains through a circular head-to-tail packing, where helix-I remains within the interior cavity. Another measure for the similarity of a given configuration to the start configuration is the fraction of native contacts, defined as^31^

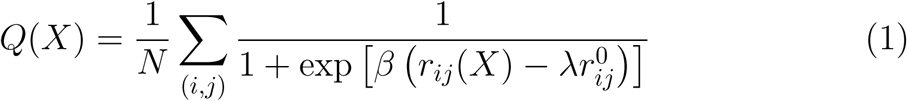

The sum runs over the *N* pairs of (*i, j*) non-hydrogen atoms *i* and *j* belonging to residues *θ_i_* and *θ_j_* that form a contact in the start configuration, i.e., their distance is less than 4.5 Å in the start configuration. Note that | *θ_i_* − *θ_j_* | *>* 3. *r_ij_*(*X*) denotes the distance between the atoms *i* and *j* in conformation *X*, while *r*_*ij*_^0^ represents that distance in the native state. *β* is a smoothing parameter taken to be 5 Å^−1^ and the factor *λ* accounts for the fluctuation when contact is formed, taken to be 1.8.

Correlations between contacts, defined here by the condition that the distance between two residues *i* and *j* (with |*i* − *j*| *>* 3) is less than 4.5 Å, are quantified by the intermittent time correlation function (TCF), *C_contact_*(*t*), which is defined as^32–34^

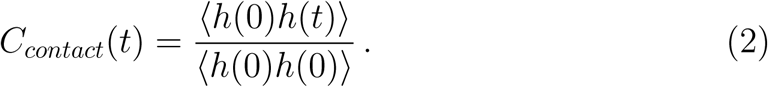

Here, *h*(*t*), a population variable, is set to one if a pair of residues is in contact at a particular time *t*, and zero, otherwise. We have also calculated similar time correlation functions for the helicity, *C_H_*(*t*), where the population variable *h*(*t*) is now set to one if a residue is in helix at a particular time *t*, and zero, otherwise.

We define the cross correlation function between residues *i* and *j*, *C*(*i, j*), as^35–37^

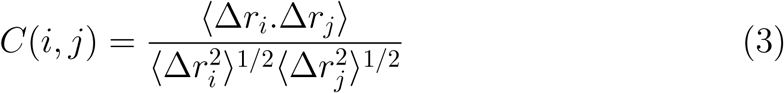

where angular brackets mark ensemble averages, Δ*r_i_* and Δ*r_j_* are the displacement vectors of the *i*-th and the *j*-th C*_α_* atoms of the protein, respectively. *C*(*i, j*) can vary by definition between +1 (complete correlated motion) and −1 (complete anti-correlated motion). Correlated residues move in the same direction, and anti-correlated residues in the opposite direction

The motion of secondary structure elements is quantified by measuring the dipole moments of the various helices, and comparing them with the ones observed in the start configurations. For this purpose, we define the dipole moment of a helix, *µ*, as

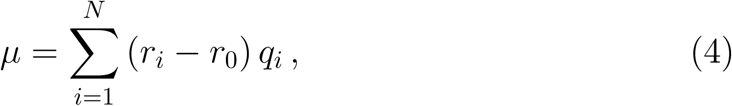

where, *r_i_* and *r*_0_ represent the position vectors of the *i*-th backbone atom and the center of mass of the helix, respectively, while *q_i_* is the partial charge of the respective backbone atom. Since the three helices differ in their number of residues, we have normalized the magnitude of the dipole moment vector by dividing it by the respective number of residues.

The stability of configurations is also evaluated by calculating the NMR N–H bond order parameter. Following Zhang and Brüschweiler,^38^ we define the order parameter for the *i*-th residue, *S*^2^ as

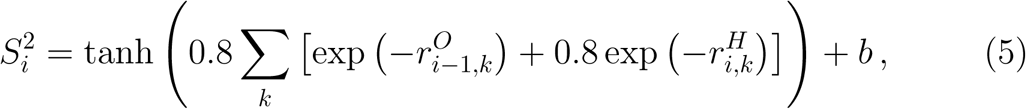

where *k* runs over all the non-hydrogen atoms except those from residues *i* and 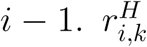 and 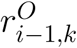 denote the distance from the non-hydrogen atom *k* to the amide hydrogen in residue *i* and carbonyl oxygen in residue *i* − 1, respectively. The parameter value *b* = −0.1 is motivated by the observation that one finds usually for rigid protein regions an order parameter value of around 0.9. Note that we have used the corrected version of *S_i_*^2,39^ according to which the distances 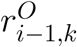 and 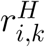 in eq 5 should be shortened by 1.2 Å.

## Results and discussion

### Hexamer

Recent experiments have established that the 104-residue serum amyloid A SAA_1−104_ assembles as a hexamer in its biologically active state, and forms in blood serum a complex with high-density lipoprotein (HDL). Dissociation of the hexamer, which is not amyloidogenic,^40^ is assisted by binding to glycosaminoglycans (GAGs), such as heparin/heparan sulfate (HS). The so-formed SAA monomers are in a second step cleaved by enzymes into shorter fragments, with the 76-residue fragment SAA_1−76_ the most commonly found species. In principle, one can think of two reasons for the cleavage. First, cleavage may aid down-regulation of SAA activity by shifting the equilibrium toward monomers, making a re-assembly toward the biologically active hexamers less likely for SAA_1−76_ than for the full-sized SAA_1−104_. A second possibility is that the shorter fragments allow for a faster degradation, allowing in this way for rapid reduction of SAA concentration.

In the present section we focus on the role of the cleavage for the equilibrium between hexamer and monomer, exploring how SAA_1−104_ and SAA_1−76_ hexamers differ in their stability and decay dynamics. In order to ensure convergence, we have monitored the time evolution of the root mean square deviations (RMSD) of the hexamer SAA_1−76_ and SAA_1−104_ with respect to their start configurations, taking into account all non-hydrogen atoms in residues 1 to 76. This choice allows us to compare RMSD values for the two systems despite their unequal length. Our results are depicted in Fig. 2 and demonstrate that both systems converge after approximately 200 ns. For this reason, we consider for further analysis only the last 300 ns of the 500 ns long trajectories. In order to test whether the elevated RMSD values of the SAA_1−76_ hexamer in relation to the SAA_1−104_ hexamer are indeed markers for differing thermodynamic stability, we have performed additional molecular dynamics simulations of the two hexamers at elevated temperatures of 325, 350, 375, and 400 K. Set-up and simulation protocol are analog to the ones described in Materials and methods, and trajectories are followed over 350 ns. The corresponding time-evolutions of RMSD are displayed in Fig. 2(c) and (d). As expected, the RMSD values rise in both systems faster and higher with increasing temperature. However, while in the case of SAA_1−104_, the final RMSD values vary within 3–4.5 Å for temperatures up to 375 K, a value of 5 Å is already reached at 325 K for SAA_1−76_. Both this difference in final RMSD values and its growth rate with time indicate a lower thermal stability of the SAA_1−76_ hexamer compared to the SAA_1−104_ hexamer.

**Figure 2:**
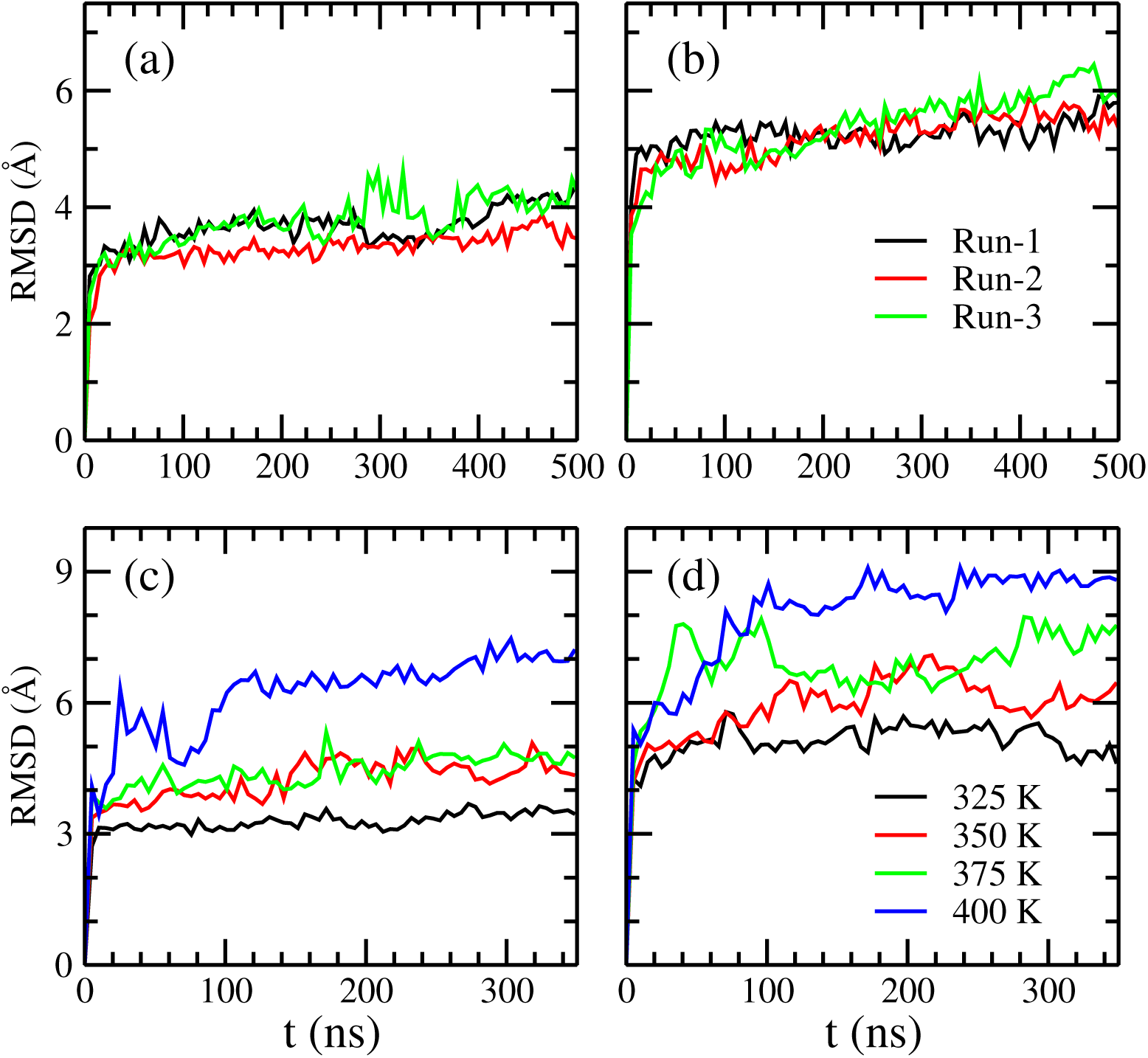
Time evolutions of root mean square deviation (RMSD) for all three trajectories of (a) SAA_1*−*104_ and (b) SAA_1*−*76_ hexamers. RMSD values are calculated with respect to the start hexamer configuration considering all the non-hydrogen atoms in residues 1 to 76 in each of the six chains. The average RMSD values as function of time for different temperatures are shown in (c) for SAA_1*−*104_ and (d) SAA_1*−*76_ hexamers.

In order to find the origin of the lower stability of the SAA_1−76_ hexamer, we have calculated the average cavity diameter <*d_cavity_*> and the solvent accessible surface area SASA of both hexamers. While the cavity diameter is similar (24.5±0.2 Å for SAA_1−76_ and 24.3±0.7 Å for SAA_1−104_), individual residues are more exposed to the solvent for SAA_1−76_, leading to the larger SASA values (55.5 (0.3) Å compared to 51.2 (0.4) Å^2^ for SAA_1−104_, see Supplemental Table SF1). This higher solvent exposure suggests that the lower SAA_1−76_ hexamer stability, leading to the higher RMSD values seen in Fig. 2, is caused by increased flexibility of individual residues in the six chains.

This higher flexibility of residues in the SAA_1−76_ hexamer results from the missing favorable inter-chain hydrogen bonds, salt-bridges, and hydrophobic interactions that stabilize the SAA_1−104_ crystal structure.^7^ This can be seen in Fig. 3 where we show in (a) the fraction of all native contacts and in (b) the same quantity restricted to *intra-chain* native contacts. Corresponding to the increase in RMSD the fraction of native contacts decreases with time for both hexamers, with the loss of native contacts more pronounced for the SAA_1−76_ hexamer than for the SAA_1−104_ hexamer. As in both cases only contacts formed by the first 76 residues are considered, it follows that the higher stability of the SAA_1−104_ hexamer is not caused by the additional contacts formed by the C-terminal tail.

**Figure 3:**
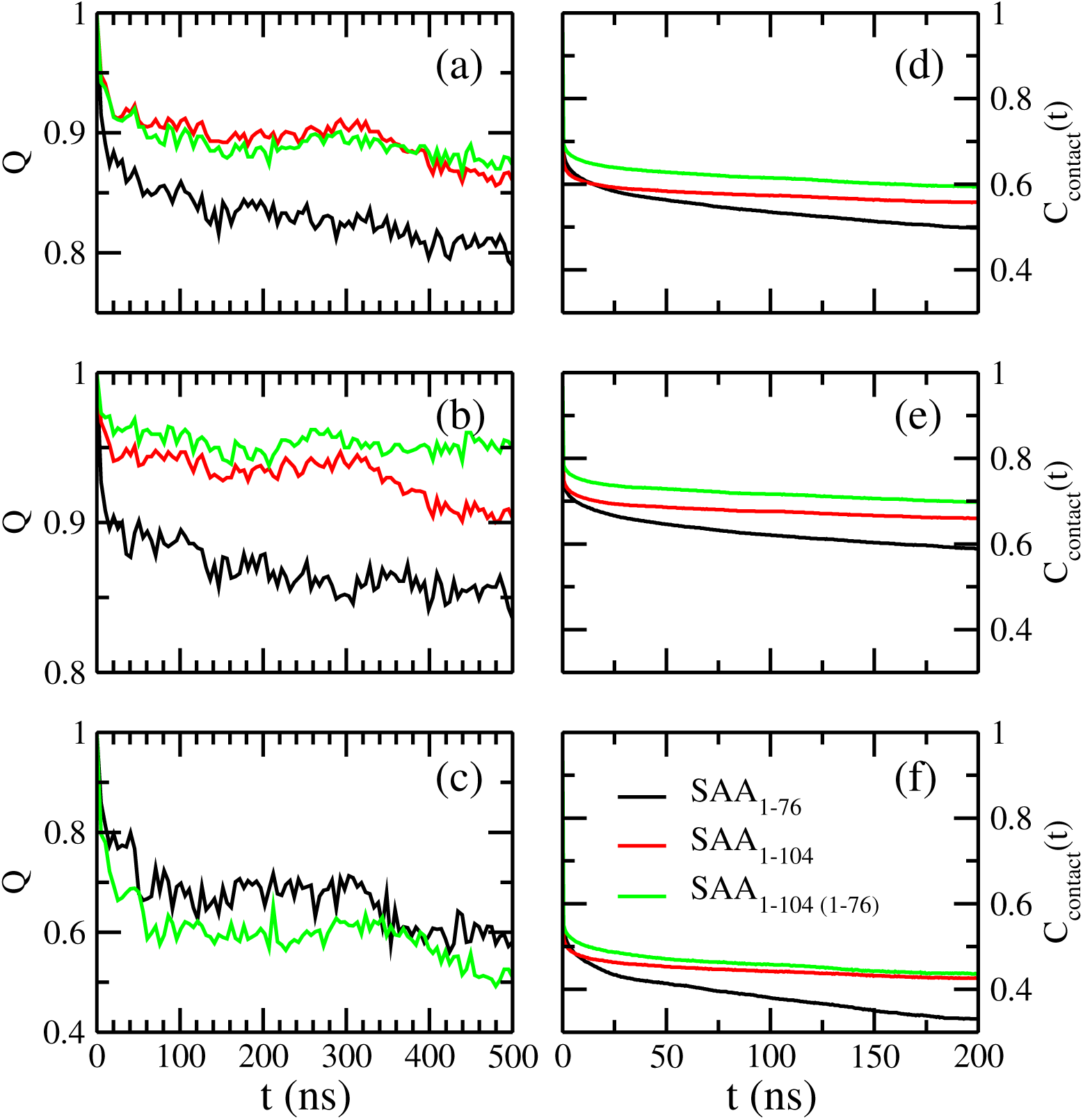
Time evolutions of the fraction of native contacts (*Q*) are shown in the left column, considering either (a) all native contacts, (b) only intra-chain native contacts, or (c) only inter-chain native contact. Data are from the first trajectory of either the SAA_1*−*76_ or the SAA_1*−*104_ hexamer simulation. The right column presents plots of the Intermittent contact time correlation function, *C_contact_*(*t*), considering either (d) all contacts, (e) only intra-chain contacts, or (f) only inter-chain contacts, formed in SAA_1*−*76_ and SAA_1*−*104_ hexamers. Data are from all three trajectories for each system, but only the last 300 ns is considered for calculating *C_contact_*(*t*). For comparison, we show also for the full-sized SAA_1*−*104_ hexamer the corresponding values for the case when only contacts between residues 1 and 76 are taken into account.

Surprisingly, the fraction of inter-chain contacts is higher for SAA_1−76_. This on first look unexpected result is likely caused by the higher flexibility of the SAA_1−76_ chains which allow them to form more easily inter-chain contacts. However, these inter-chain contacts are transient and do not contribute to the stability of the hexamer. This can be seen from Fig. 3(d-f) where we show time correlation functions (TCFs) of the contacts, taking into account either all contacts, or considering only either inter-chain or intra-chain contacts. Irrespective of the types of contacts, the TCFs for the SAA_1−76_ hexamer decay fast and monotonically, while for SAA_1−104_ hexamer the decay is slower and quickly approaches a plateau. This is also the case when for the later only residues 1-76 are considered. The decay time is especially short for the inter-chain contacts in the SAA_1−76_ hexamer, demonstrating the short life times and transient nature of these contacts. Hence, the additional inter-chain contacts can only partially compensate for the loss of stability resulting from the reduced number of intra-chain contacts, as their life times are short, and overall the total fraction of native contacts is lower for SAA_1−76_ hexamers than for SAA_1−104_ hexamers. Hence, the intra-chain contacts rather than the inter-chain contacts determine the overall stability of the SAA hexamers.

The above discussion implies that when part of the hexamer, the intrinsic stability of the folded SAA_1−104_ chains is higher than for the SAA_1−76_ chains. This is supported by Fig. 4(a) where we show the residue-wise NMR order parameter (*S*^2^) of the first 76 residues of both proteins, averaged over all six chains in a hexamer and all three trajectories. The lower the value of *S*^2^, the higher will be the flexibility of the respective N–H bond and hence the corresponding residue and vice versa. Comparing the two proteins, we find a signal for increased flexibility of SAA_1−76_ chains in two regions, one given by residues 30–43, and the other made of residues 63–76. These residues have a higher probability to interact with the solvent, starting in this way the dissociation process of the hexamer. Fig. 4(b) confirms indeed that the solvent accessible surface area (SASA) of these residues, belonging to either helix-II or helix-III, is higher for SAA_1−76_ than for the full-sized SAA_1−104_ where the C-terminal tail (including helix-IV) protects these residues from being exposed to the solvent.

**Figure 4:**
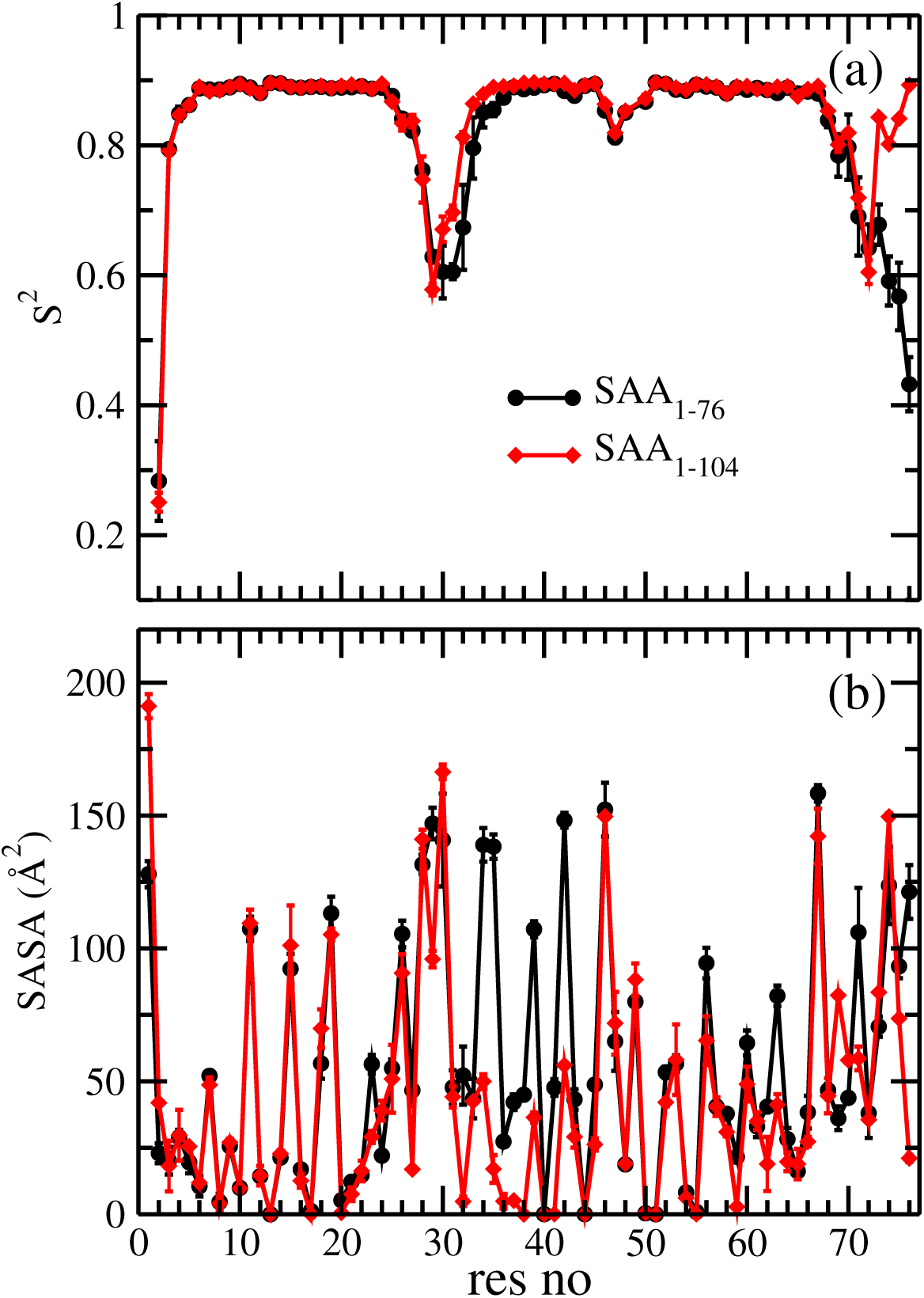
Residue-wise (a) backbone N–H order parameter (*S*^2^) and (b) solvent accessible surface area (SASA) for the SAA_1*−*76_ and SAA_1*−*104_ hexamers as calculated over the last 300 ns of all three trajectories. Data are shown only for the first 76 residues.

### Monomer

In the above section we have demonstrated that hexamers formed by SAA_1−104_ have a higher stability than such formed from the fragment SAA_1−76_. Hence, it is unlikely that SAA_1−76_ and similar fragments, once generated by enzymatic cleavage from the full sized protein, will re-assemble into a hexamer; and if formed by chance, such hexamer would decay quickly. However, it is not clear whether the sole purpose of the cleavage is inhibiting re-assembly into the biologically active hexamer. Another reason for the cleavage could be to decrease the SAA concentration by easing degradation of the protein. We have already shown in the previous section that within the hexamer the SAA_1−76_ chains have lower internal stability than full-length SAA_1−104_ chains. A similar increased flexibility of the isolated chains may allow for the fragments structural changes that could encourage proteolysis. However, the higher flexibility would also increase the risk of aggregation as it could lead to release of the first eleven residues from helix-I. These N-terminal residues are considered to be crucial for amyloid formation. Hence, in order to probe the effect of the cleavage on stability and potential amyloid-forming tendencies of SAA monomers, we have studied also the isolated monomers of the full-length SAA_1−104_ and the SAA_1−76_ fragment in another set of molecular dynamic simulations.

Similar to the hexamer, we first establish what part of the simulated trajectories can be used for analysis. For this purpose, we monitor for the two monomers the time evolution of RMSD with respect to their start configurations, considering again only the non-hydrogen atoms in the first 76 residues. Our results, shown in Supplemental Fig. SF1, are similar to the hexamers in that the changes of RMSD are smaller for SAA_1−104_ monomers (around 2 Å) than for the SAA_1−76_ monomers (around 6-12 Å) which have again more pronounced fluctuations. However, as a plateau is approached for both molecules, and the simulations converge in about 500 ns, 500 ns of the 1 *µ*s-long trajectories remain for our analysis.

The stability differences between SAA_1−76_ and SAA_1−104_ monomers, seen in the time evolution of RMSD, are further investigated by comparing the average number of native contacts (<*N_nat_*>), residue-residue contacts (<*N_c_*>), and helicity (<*h*>). In Table 1, we list the differences of these values to the ones measured in the respective start configurations. For the full-sized protein, we have also calculated these values restricted to the first 76 residues, and have added the values in the table to allow for a better comparison with the fragment SAA_1−76_. The twice smaller values of <*N_nat_*> for SAA_1−76_ monomers (in relation to full-length monomers) confirm the larger deviations, and in turn lower stability, of the truncated fragment. This difference is also seen when only the first 76 residues of the full-sized protein are considered. Similarly, the loss of helicity <*h*> is larger in the fragment than in the full sized protein.

**Table 1:**
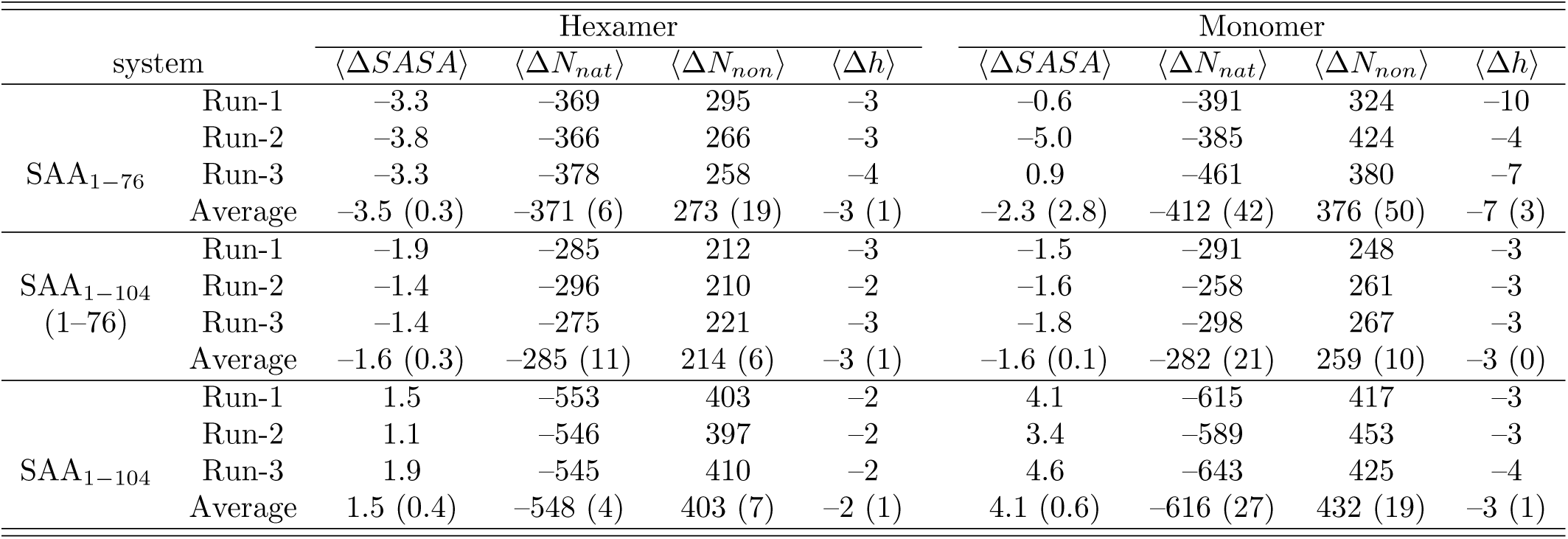
Difference of various quantities with respect to the start configuration, measured in all three trajectories of SAA_1*−*76_ and SAA_1*−*104_ chains in the hexamer and in isolated monomers. We list differences calculated for the solvent accessible surface area <Δ*SASA*> per residue (in Å^2^), number of native contacts (<Δ*N_nat_*>), number of non-native contacts (Δ*N_non_*), and helicity (Δ*h*). For comparison we present also data for SAA_1*−*104_ chains considering only the first 76 residues. Averages are taken over the three trajectories with the standard deviations listed in parenthesis.

|  |  | Hexamer |  |  |  | Monomer |  |  |  |
| --- | --- | --- | --- | --- | --- | --- | --- | --- | --- |
| system | | $\langle\Delta SASA\rangle$ | $\langle\Delta N_{nat}\rangle$ | $\langle\Delta N_{non}\rangle$ | $\langle\Delta h\rangle$ | $\langle\Delta SASA\rangle$ | $\langle\Delta N_{nat}\rangle$ | $\langle\Delta N_{non}\rangle$ | $\langle\Delta h\rangle$ |
| SAA<sub>1-76</sub> | Run-1 | -3.3 | -369 | 295 | -3 | -0.6 | -391 | 324 | -10 |
|  | Run-2 | -3.8 | -366 | 266 | -3 | -5.0 | -385 | 424 | -4 |
|  | Run-3 | -3.3 | -378 | 258 | -4 | 0.9 | -461 | 380 | -7 |
|  | Average | -3.5 (0.3) | -371 (6) | 273 (19) | -3 (1) | -2.3 (2.8) | -412 (42) | 376 (50) | -7 (3) |
| SAA<sub>1-104</sub><br>(1-76) | Run-1 | -1.9 | -285 | 212 | -3 | -1.5 | -291 | 248 | -3 |
|  | Run-2 | -1.4 | -296 | 210 | -2 | -1.6 | -258 | 261 | -3 |
|  | Run-3 | -1.4 | -275 | 221 | -3 | -1.8 | -298 | 267 | -3 |
|  | Average | -1.6 (0.3) | -285 (11) | 214 (6) | -3 (1) | -1.6 (0.1) | -282 (21) | 259 (10) | -3 (0) |
| SAA<sub>1-104</sub> | Run-1 | 1.5 | -553 | 403 | -2 | 4.1 | -615 | 417 | -3 |
|  | Run-2 | 1.1 | -546 | 397 | -2 | 3.4 | -589 | 453 | -3 |
|  | Run-3 | 1.9 | -545 | 410 | -2 | 4.6 | -643 | 425 | -4 |
|  | Average | 1.5 (0.4) | -548 (4) | 403 (7) | -2 (1) | 4.1 (0.6) | -616 (27) | 432 (19) | -3 (1) |

On the other hand, the total number of contacts <*N_c_*> is approximately the same in the truncated fragment and between residues 1-76 of the full-sized protein, indicating that the fragment not only loses helical contacts but that the structure of the fragment changes and new contacts are formed.

For the full-sized protein, the loss of native contacts is mainly in the C-terminal region, with only few (about 10 %) of native contacts within residues 1-76 lost in the first 1*µ*s. This demonstrates the protective role of the C-terminal segment of residues 77-104, and indicates that after cleavage, the fragment SAA_1−76_ quickly loses its tertiary structure. This picture is supported by the side-chain–side-chain contact maps shown in Supplemental Fig. SF2, which shows that absence of the C-terminal tail leads in the SAA_1−76_ to loss of contacts between residues 70–76 and 60–69, i.e., the cleavage increases the flexibility of helix-III. In turn, contacts between residues 32–38 and 65–67, located on helix-II and helix-III, are also dissolved. A similar loss of inter-helix contacts is seen between helix-I (residues 26-28) and helix-II (residues 32-34). This reduction in the number of contacts increases the residue-wise solvent accessible surface area in the fragment, see also Supplemental Fig. SF3. Similar to that of the hexamer, most of the residues present in helix-II of the truncated SAA monomer become more exposed toward the solvent. The difference in SASA between fragment and full-sized monomer is largest for the residues 30-40, belonging to helix-II, and about 10% larger in the monomer than in the hexamer. This SASA difference results from the weaker contacts of helix-II with the adjunct helices I and III in the fragment. This is consistent with the fact that, as shown in Supplemental Fig. SF2, helix-II loses the contacts with both helix-I and helix-III that are present in the native structure. However, the SASA values of the residues 57, 58, 61, 64, 65, 68, and 69 on helix-III, which in the native structure form contacts with helix-II, do not increase after losing the contacts. Instead, the SASA values even decrease, indicating that these hydrophobic residues have formed alternative contacts, leading to a structural re-arrangement of the remaining three helices.

Hence, cleavage of the C-terminal tail destabilizes the native structure of SAA monomers through reducing inter-helix contacts and increased solvent exposure. As soon as (after release from the hexamer) the full-length SAA monomers are cleaved and lose the C-terminal tail (including helix-IV), their native structure decays, and the chains are unlikely to re-associate into a hexamer. On the other hand, unfolding of the full-length monomer structure (SAA_1−104_) happens, if at all, only on much longer time scales. These longer life times allow the full-length protein, unlike the fragment, to re-assemble into a hexamer.

However, the reduced stability of the fragment SAA_1−76_, affects also the N-terminal eleven residues, which are known to be critical for SAA amyloid formation. The strength of this effect depends on the length of the cleaved fragment. This can be seen in Fig. 5 where we show for SAA fragments of various length the residue-wise NMR order parameter (*S*^2^) for the backbone N–H bond vectors (eq. 5). The flexibility of the first eleven N-terminal residues increases drastically with subsequent cleavage of the helices, facilitating evermore misfolding into an aggregation prone configuration. A special role here seems to fall to the segment SAA_1−76_ which is in between the two extreme cases of the N-terminal segment fully cached in helix-I (the full-sized protein made of all four helices), or of the N-terminal segment being highly flexible and stabilized only by the environment of helix-I (SAA_1−27_) and helix-II (SAA_1−47_). This intermediate position of the SAA_1−76_ fragment therefore indicates that helix-III takes a special role in the misfolding and aggregation of SAA.

**Figure 5:**
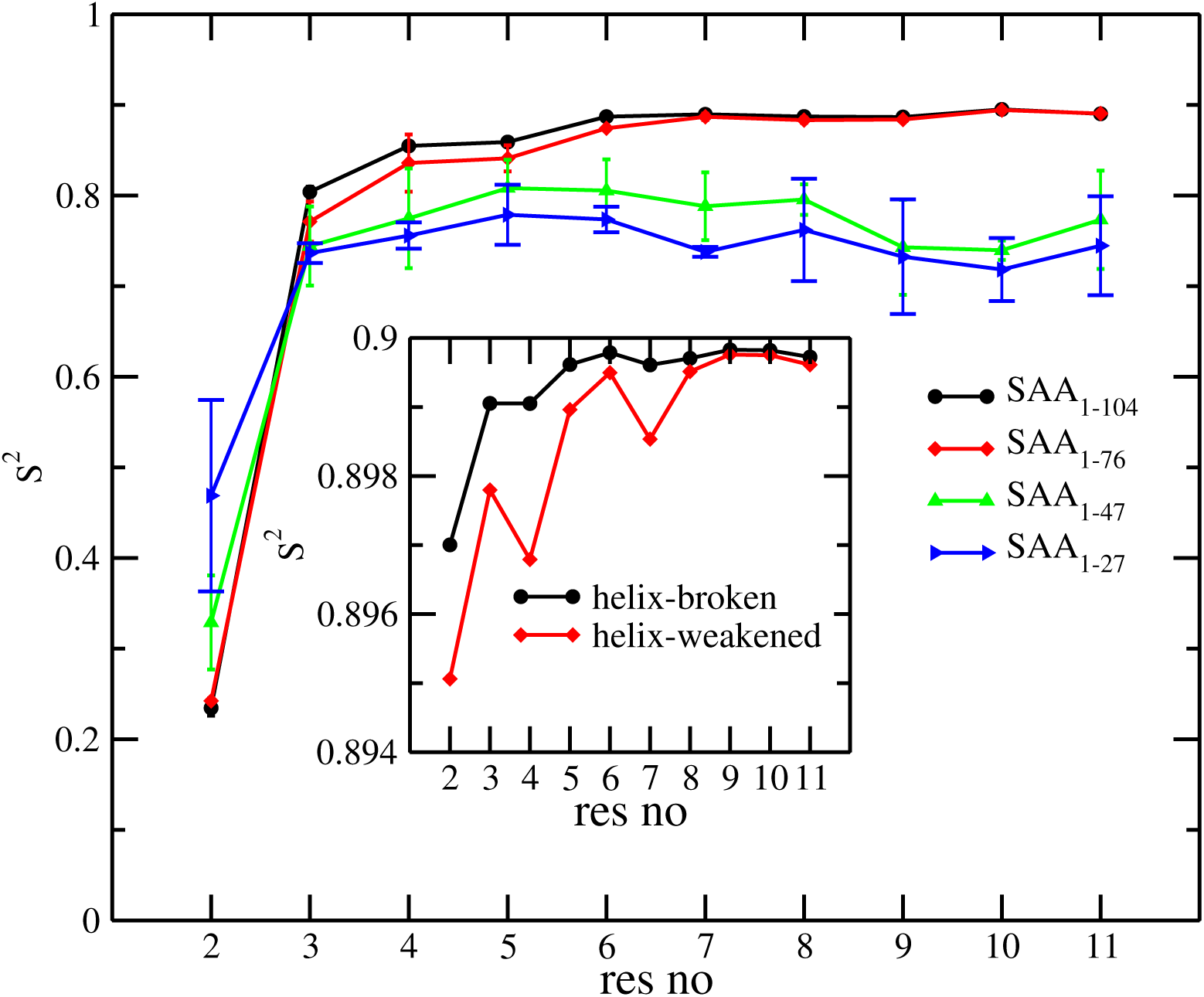
Residue-wise backbone N–H order parameter (*S*^2^) for the first eleven residues of different SAA protein monomer fragments, (1-104), (1–76), (1–47), and (1–27), as calculated over the last 500 ns of all three trajectory of each system. For the fragment SAA_1*−*76_ we distinguish further between helix-broken and helix-weakened structures (see text). Residue-wise backbone N–H order parameter (*S*^2^) for the first eleven residues corresponding to these two structures of SAA_1*−*76_ monomers is shown in the inset.

In order to understand this special role of helix-III, we have calculated for each helix the corresponding dipole moment. The time evolutions of the per-residue dipole moment (*µ*) for all three helices are displayed in Fig. 6(a-c) where we depict for this quantity the magnitude of the difference between actual value and the one measured in the start configuration. While the dipole moments of helix-I and helix-II change little with time, i.e., are not affected by the cleavage of the C-terminal residues, there is a clear signal for helix-III. The here observed change in dipole moment could indicate either breakage of helix-III into two shorter segments connected by a kink (named by us a helix-broken configuration), or a shortening of the helix-III with the released residues taken random orientations (called by us a helix-weakened configuration). Visual inspection shows that both possibilities happen, the helix-broken case in run 2, and the helix-weakened one in run 1 and 3. The relative orientation of helix-III with respect to the other two helices changes in both motifs, as can be seen from the relative orientation between pairs of helices in Figs. 6(d-f) and the center-of mass distances between these pairs in Figs. 6(g-i). This is particularly noticeable for the helix-broken case (Figs. 6(e) and (h)).

**Figure 6:**
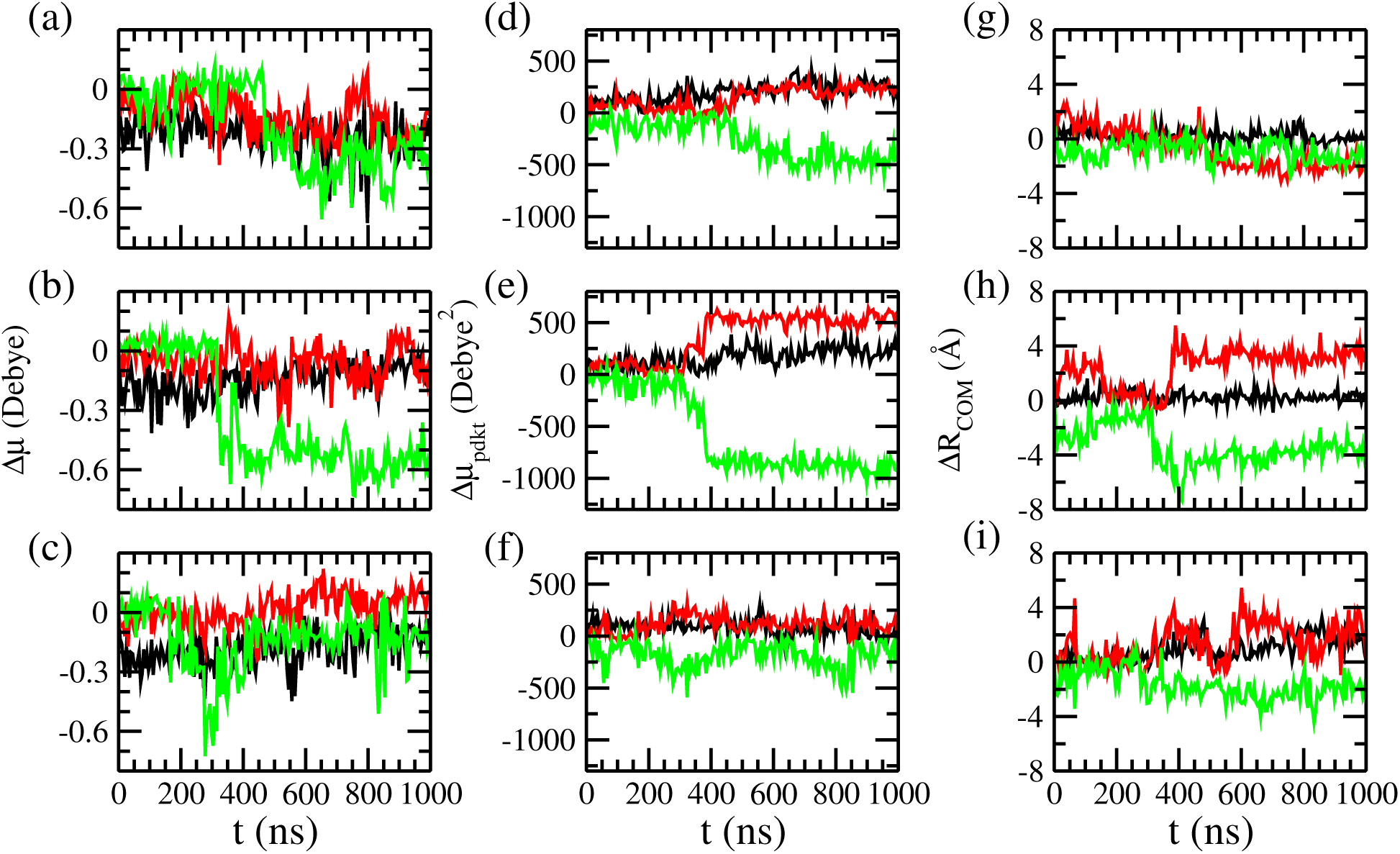
Time evolution of (a-c): difference in dipole moments (*µ*) of a helix at a given time minus the corresponding value in the start configuration. Shown are the magnitude of this difference divided by the number of residues for helix-I, helix-II, and helix-III in the SAA_1*−*76_ fragment. The color-code is as follows: helix-I is shown in black, helix-II in red, and helix-III in green. In (d-f) we show the scalar product of dipole moment vectors and in (g-i) center of mass distances between pairs of helices. The color-coding is as follows: values for helix pair (I,II) are displayed in black, for helix pair (II,III) in red, and for helix pair (I,III) in green. The results for run-1 is shown in the first row, for run-2 in the middle row, and for run-3 in the last row.

In order to go beyond visual inspection, we introduce the following definition for the two motifs. In a helix-weakened configurations, helix-III is still preserved for residues 50–62, but at most three residues are still helical in the C-terminus of helix-III (residues 63–69). On the other hand, a kink is formed within residues 50 to 62 in case of helix-broken configuration (i.e., the helicity of this segment is less than 13) and the C-terminus of helix-III is preserved (at least five residues are helical in the segment 63–69). In both cases, the orientation of helix-III changes toward helix-I, with the C*_α_* residue distance between residue 1 and 69 (d*C_α_*) less than 20 Å in case of helix-broken configuration, and more than 20 Å in the helix-weakened case.

Our above results show that upon cleavage of the C-terminal residues 77-104, the position of helix-III is no longer restrained by contacts with these residues, but can now form contacts with helix-I. If helix-III moves toward the C-terminal end of helix I, the lack of helix-stabilizing contacts will lead to a shortening of the helix and a disordered segment at the C-terminus becomes oriented toward the C-terminus of helix-I. On the other hand, if helix-III moves toward the N-terminus of helix-I, helix-III breaks up into two shorter pieces connected by a kink around the residues 55–58, with the C-terminal end of helix-III now pointing toward the first eleven N-terminal residues of helix-I. We believe that this re-alignment of helix-I and helix-III in the two motifs is correlated with a possible release of the N-terminal residues 1-11 from helix I, allowing these residues to re-fold into a *β*-hairpin as needed for attachment of other chains and starting amyloid formation. This assumption is supported by Fig. 7 where we show how rapidly contact number and helicity, measured for the first eleven N-terminal residues, change with time. This speed of change is quantified by the intermittent time correlation function (TCF) of the two quantities, calculated for the helix-weakened case from the last 500 ns of run 1 and 3, and the helix-broken case from the last 500 ns of run 2. For comparison, we show values obtained for the full-sized SAA_1−104_ monomer. While for the helix-weakened configuration both the helicity and the number of contacts of the N-terminal segment decay faster than in the more stable full-sized SAA_1−104_ monomer, there is no qualitative difference seen between the helix-broken configuration and the full-sized protein. Hence, our results suggest a higher chance for the release of the first eleven N-terminal residues from helix-I in helix-weakened configurations than in helix-broken configurations, where the segment is stabilized by contacts with the C-terminus of helix-III.

**Figure 7:**
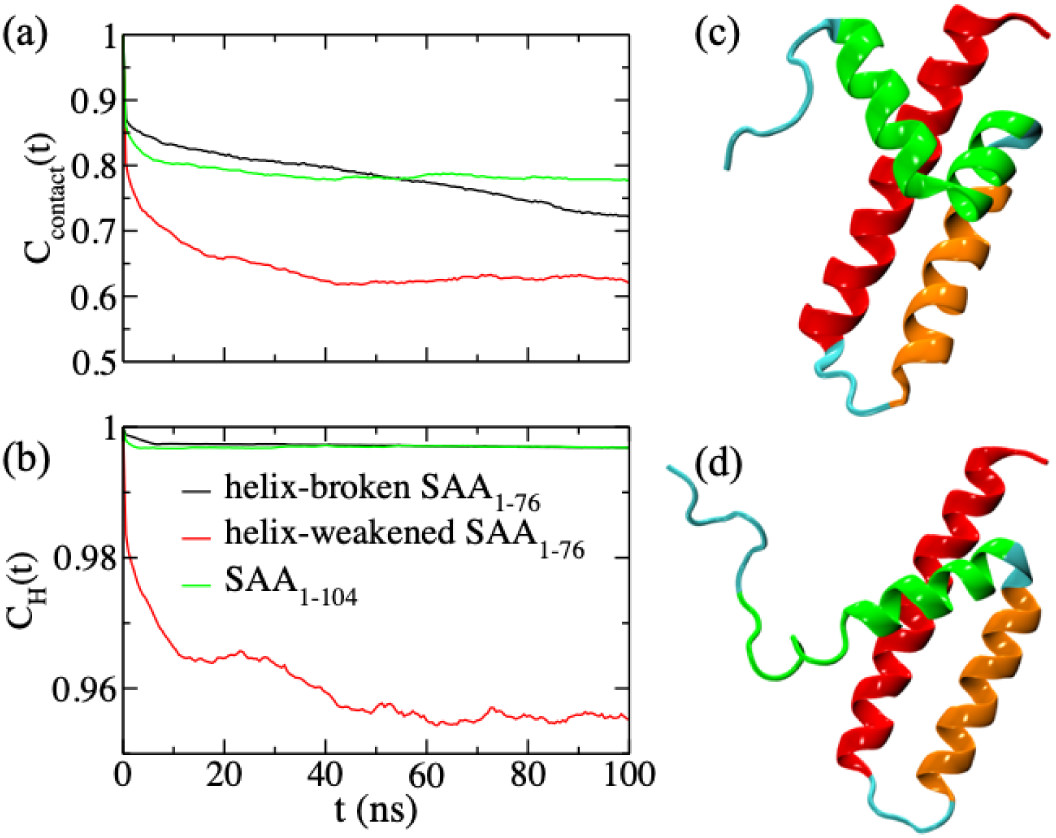
Intermittent time correlation function for (a) the contact *C_contact_*(*t*) and (b) the helicity *C_H_* (*t*), where contacts and helicity are calculated for the first eleven N-terminal residues of helix-I of either helix-broken or helix-weakened SAA_1*−*76_ conformations. As a reference, data for SAA_1*−*104_ monomers are also shown. Representative configurations for the (c) helix-broken and (d) helix-weakened SAA_1*−*76_ structures are presented in the right subfigures. The color scheme of the three helices, helix-I to III, is same as in Fig. 1

In order to probe in more detail how the relative movement of helix-III in the two motifs affects residues in other parts of the molecule, especially the first eleven N-terminal residues, we show in Figs. 8(a-b) and (c-d), respectively, for both motifs the contact map and cross correlation function, *C*(*i, j*), between the residues. In the helix-weakened structure the C-terminal residues of helix-III form contacts with the C-terminal residues of helix-I. Formation of these contacts is by a correlated motion of C-terminal of helix-III (residues 58–69) with the C-terminal of helix-I (residues 19–27), and a consequently anti-correlated motion with N-terminal of helix-I (residues 1– 11), driving them in opposite directions. This motion is reversed in the helix-broken structure, where these contacts are absent and instead the C-terminus of helix-III forms contacts with the N-terminal residues of helix-I (here primarily the first eleven residues). These additional contacts, that do not exist in the helix-weakened structure, stabilize helix-I in the helix-broken structure, i.e., prevent release of the first eleven residues and their misfolding into an amyloid-prone configuration. The motif is further stabilized by a re-orientation of the hydrophilic residues on helix-I, which in the native structure point toward helix-II, but in the helix-broken motif face outward to the solvent. On the other hand, in the helix-weakened motif, the hydrophilic residues stay oriented toward helix-II, but the hydrophobic residues now face toward the solvent, which in the native structure is the case for only 30% of the hydrophobic residues on helix-I. Note that exposure of such large hydrophobic patches as seen on helix-I in the helix-weakened structure often serves as a signal for activating degradation by the proteosome.

The already existing stable contacts between the C-terminal end of helix-I and helix-II in the helix-weakened structure restrict the possibility of formation of new contacts between the C-terminal end of helix-III and the C-terminal end of helix-I. This also simultaneously restricts the formation of salt-bridges between the opposite charged residues or hydrogen bonds between the side chains. On the contrary, repulsion between the similar charged residues on the C-terminal ends of helix-III and helix-I will further destabilize the overall structure. The only reasonably stable contacts that we found in helix-weakened structures are Y21–R62, M24–I65, and R25–I65. However, these contacts are neither sufficient to compensate the charge-charge repulsion, and the helix-weakened structure is flexible, with frequent transitions toward the original arrangement of helix-I and helix-III. The net-effect is a loss of the forces that stabilize helix-I in the native structure of the full-sized SAA_1−104_ monomer, but are missing in the shorter segments. The resulting higher entropy of helix-I leads to its de-stabilization, especially at the N-terminus as the C-terminus is still supported by contacts with helix-III. As a consequence, there is an increased chance for the first eleven residues to be released from helix-I and to misfold into strand-like configuration (see inset of Fig. 5) that may start the aggregation process by way of a fly-fishing mechanism as suggested by G. Reddy (private communication).

On the other hand, in helix-broken structures, strong contacts are formed between residues F4–F68, L7–I65, and F11–F69. The phenyl rings of the pair of Phe residues i.e., (F4, F68), and (F11, F69) orient in such a way that they remain one upon another with an inter-planar angle of ∼ 0^◦^, resulting in strong *π*–*π* stacking interactions of the phenyl rings. The result is a helix-helix linkage between the C-terminal end of helix-III and free N-terminal end of helix-I, that not only stabilizes the overall structure but especially the N-terminus, keeping the first eleven residues cached in helix-I and therefore reducing the probability of their misfolding and subsequent aggregation.

Hence, based on our observations, we propose the following mechanism for tuning SAA concentration. After the release of SAA_1−104_ chains from the hexamer enzymatic cleavage leads to SAA_1−76_ monomers that are unstable as they lack the helix–helix linkage interaction between helix-III and helix-IV. The C-terminal of helix-III becomes exposed, and helix-III can now move relative to helix-I. This process is facilitated by formation of transient contacts between residues 20–21 (in helix-I) and 61–62 (in helix-III), which now permit movement of the C-terminus of helix-III toward either the C-terminus or N-terminus of helix-I, i.e., enabling transitions between the two resulting structures, coined by us helix-weakened and helix-broken structures. With only three runs we do not have sufficient statistics to quantify the relative frequency of the two motifs, but our simulations indicate that after cleavage of the full-length protein, most SAA_1−76_ monomers evolve into a helix-weakened configuration.

We conjecture that in helix-weakened configurations the large exposed hydrophobic patches on helix-I will activate further proteolytic degradation, likely by protease cathepsin D^41^ after binding with the heat shock protein Hsp70^41–43^ recruited in response to the primary disease. The net-effect would be a reduction in SAA concentration. However, the larger flexibility of the N-terminal segment of the first eleven residues also increases the risk that that these residues unfold from helix-I and take strand-like configurations which could nucleate amyloid formation. Under neutral pH the helical conformation of the segment is stabilized through a transient salt bridge between residues 1R and 9E. This salt bridge (defined by us by the condition that the center-of-mass distance between the ammonium or carboxylate groups of residues 1R and E9 is below 4Å) is formed in about 70% of the full-length SAA_1−104_ monomers, but only in about 25% of the helix-weakened SAA_1−76_ configurations. On the other hand, in helix-broken configurations, this salt bridge is found with similar frequency as in the full length protein, and the N-terminal segment is stabilized additionally by interactions with the C-terminal of helix-III. Hence, the helix-broken configurations are less aggregation-prone than helix-weakened configurations. We hypothesize that this difference is minor under neutral conditions, making helix-weakened configurations more desirable than helix-broken configurations which do not have large hydrophobic patches exposed to the solvent and therefore are more difficult to degrade.

The situation may be different under acidic conditions such as seen in conjunction with cancer or or inflammatory diseases^44,45^ where this salt bridge between residues 1R and 9E can no longer be formed. In preliminary simulations, designed to mimic acidic conditions, the salt bridge is found at pH=5 in only about 40% for helix-broken and about 15% for helix-weakened configurations; and at pH=4 it is less than 5% of the SAA_1−76_ configurations. The disappearance of the salt bridge will likely increase the chance for aggregation and amyloid formation, with the risk mitigated in helix-broken configurations by the additional contacts between the N-terminal segment and the C-terminus of helix-III. Interestingly, we observe in the same set of simulations also a shift from helix-weakened to helix-broken configurations when going to acidic conditions, see Fig. 8(e) and (f). This transition between the two forms therefore counteracts the increased danger of unfolding of the N-terminal segment by a structural conversion that stabilizes this segment through additional contacts with helix-III. Hence, this pH-driven transition between helix-weakened and helix-broken configurations appears to be a mechanism to counteract the increased chance for aggregation and amyloid formation otherwise often seen under acidic conditions.^44,45^

**Figure 8:**
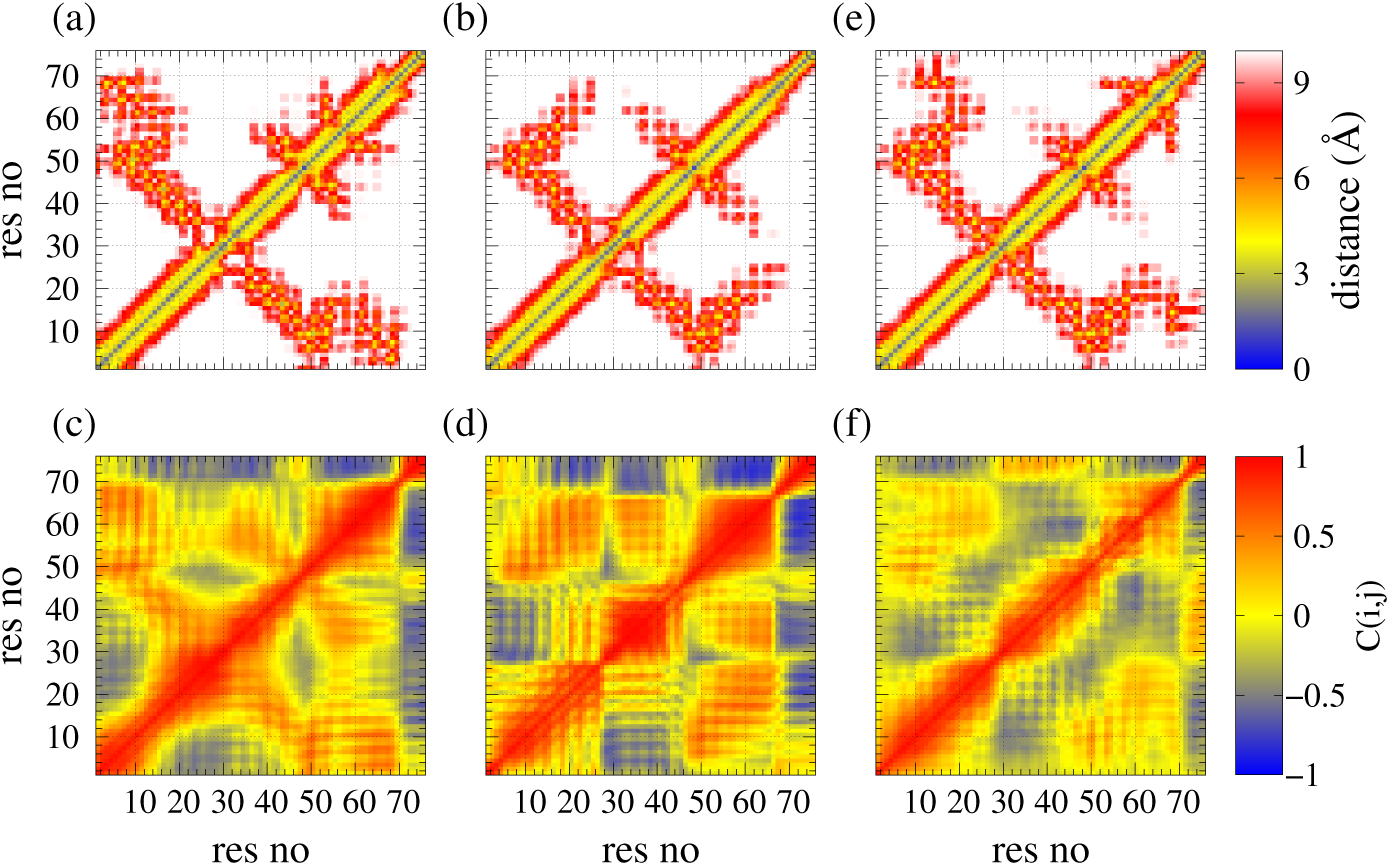
Side-chain–side-chain contact map of (a) helix-broken and (b) helix-weakened SAA_1*−*76_ monomer configurations at neutral pH, showing the distance between pairs of residues. The corresponding two-dimensional dynamic cross-correlation map *C*(*i, j*) of these pairs are shown for helix-broken configurations in (c) and for helix-weakened configurations in (d). For the later motif we also show in (e) and (f) corresponding figures obtained from simulations under acidic conditions (mimicking a pH=4).

## Conclusions

Various diseases cause over-expression of the serum amyloid A protein (SAA) leading in some, but not all, cases to amyloidosis as a secondary disease. The response to over-expression involves dissociation of the SAA hexamer and subsequent enzymatic cleavage of the full-length SAA_1−104_ chains into shorter fragments, most commonly the 76-residue fragment SAA_1−76_. Analyzing extensive molecular dynamics simulations we propose a mechanism to down-regulate SAA activity and concentration that relies on this cleavage and is sketched in Fig. 9. As serum amyloid A in its functional form assembles into hexamers, we have first tested the hypothesis that the cleavage shifts the equilibrium for SAA_1−76_ fragments from the biologically active hexamers to potentially amyloidogenic monomers. Our molecular dynamics simulations confirm that hexamers built from full-length SAA_1−104_ chains are indeed more stable than such formed from SAA_1−76_ fragments. We explain this lower stability with the larger exposure of helix-II and helix-III in the SAA_1−76_ hexamer chains. This lower stability reduces the chance for the SAA_1−76_ fragments to re-assemble into a hexamer, and if formed by chance, such hexamer would decay quickly.

**Figure 9:**
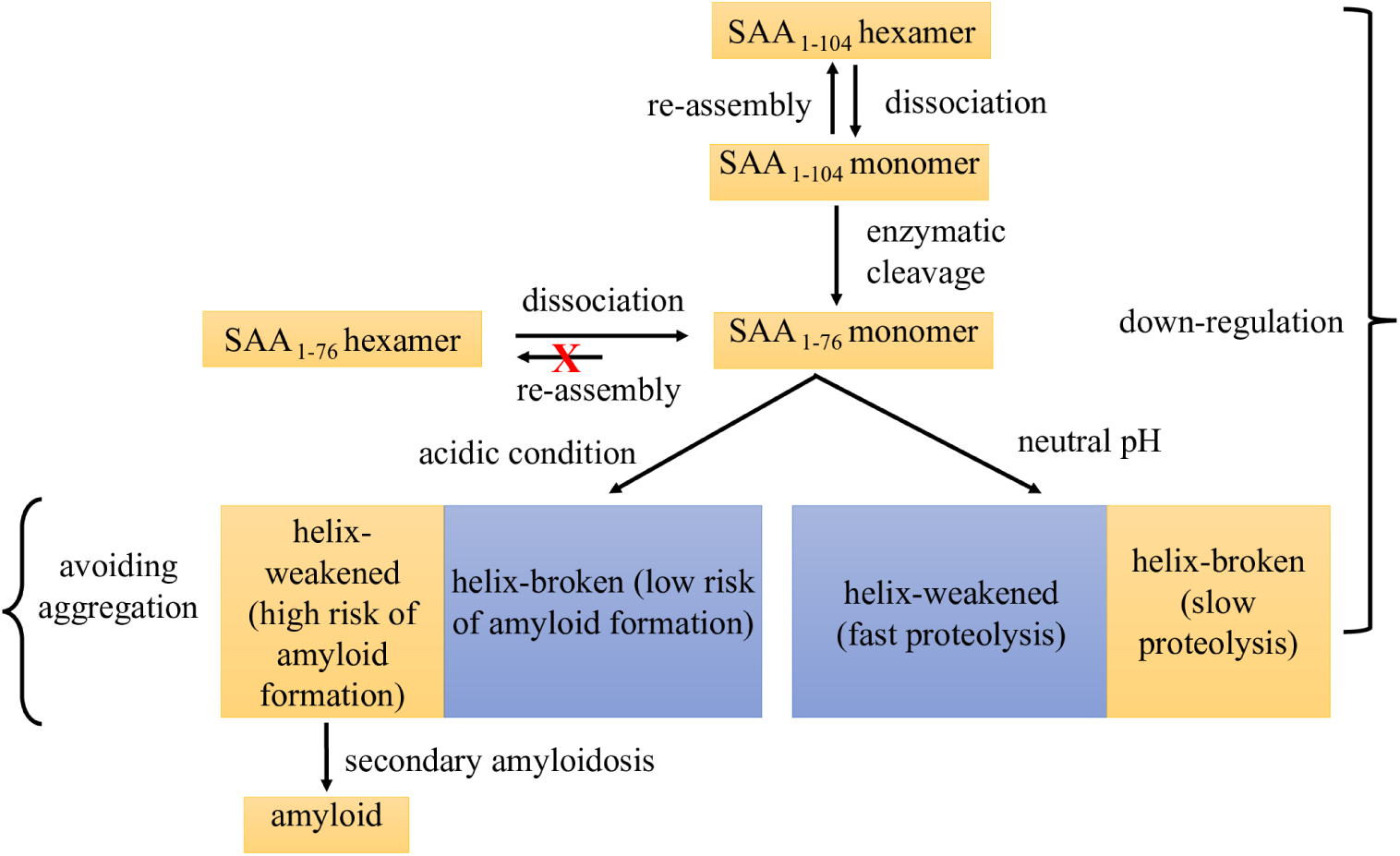
Sketch of the proposed mechanism that balances the desire for down-regulating SAA activity and concentration with the need to avoid harmful aggregation.

Our hexamer simulations also suggested a higher internal stability of the full-length SAA_1−104_ chains compared with the SAA_1−76_ fragments. As the reduced stability of the fragment likely triggers further degradation, reducing SAA concentration, we have added molecular dynamics simulations of isolated monomers to probe also the lower stability of the SAA_1−76_ fragments. As the first eleven residues are crucial for amyloid formation,^18^ we were especially interested in the question of how the cleavage affects the stability of this N-terminal segment which is part of helix-I (residues 1–27) in the native structure and protected by the C-terminal tail (including helix-IV). On the other hand, in SAA fragments too short to contain helix-III (residues 50-69), helix-I is less protected, and only a transient salt bridge between residues 1R and 9E stabilizes the helical configuration of the N-terminal segment.^19^ Especially under acidic conditions where this salt bridge cannot be formed, the N-terminal residues may unfold to form strand-like segments^19,20^ and to nucleate aggregation.

SAA_1−76_ fragments are on the cusp between structures where the N-terminal residues are firmly integrated in helix-I (as in the full-length protein), and such where they are flexible enough to be released easily from helix-I, see Fig. 5. This is because interactions with helix-III may either stabilize or de-stabilize helix-I. Since the fragment SAA_1−76_ lacks the helix–helix linkage interactions between helix-III and helix-IV, seen in the full-length protein, the C-terminus of helix-III is able to move toward either the C-terminus of helix-I or toward the N-terminus of helix-I. The first and more common case leads to helix-weakened structures characterized by a weakened helix-III causing large exposed hydrophobic patches on helix-I that will trigger further degradation by the proteosome, therefore reducing the SAA concentration and down-regulating SAA activity. However, this motif is also characterized by a reduced stability of helix-I which raises the probability for a release of the aggregation-prone first eleven residues, providing a potential start point for subsequent amyloid formation. This is especially the case under acidic conditions where the transient salt bridge 1R-9E cannot be formed that stabilizes the helical conformation of the N-terminal segment under neutral conditions. On the other hand, in the helix-broken structures, newly formed contacts between the C-terminus of helix-III and the N-terminus of helix-I stabilize the latter helix, reducing the probability that the aggregation-prone first eleven residues are released from helix I. Hence, the increased risk for aggregation and amyloid formation associated with acidic conditions is mitigated by switching from helix-weakened to helix-broken configurations, a process that we have observed in preliminary simulations designed to mimic an acidic environment. The possibility for such transitions may be the reason why cleavage of the full-length SAA protein leads most often to SAA_1−76_ fragments, which by switching between helix-weakened and helix-broken configurations can optimize the chance for degradation while minimizing the risk of aggregation and amyloid formation. In most patients where colon cancer, inflammatory bowel disease, or rheumatoid arthritis leads to over-expression of SAA, the described mechanism enables down-regulation of activity and concentration of SAA. We speculate that SAA amyloidosis indicates failure of this switching mechanism.

## Acknowledgements

The simulations in this work were done using the SCHOONER cluster of the University of Oklahoma and XSEDE resources allocated under grant MCB160005 (National Science Foundation). We acknowledge financial support from the National Institutes of Health under grant GM120634.

## Author contributions

W.W. and P.K. contributed equally in simulation and analysis. W.W., P.K., and U.H.E.H. designed this study and wrote the paper.

## Correspondence

Correspondence and requests for materials should be addressed to W.W., P.K. or U.H.E.H.

