## Supplemental figures and tables for "Cleavage, down-regulation and aggregation of serum amyloid A"

Table SF1: Solvent Accessible Surface Area ($\langle SASA \rangle$) per residue averaged over all chains in the $SAA_{1-76}$ and $SAA_{1-104}$ hexamers. We show both the values for the individual trajectories and for the resulting averages (with standard deviation listed in parenthesis). For comparison we list also the respective values for the free monomers.

| system | | Hexamer | Monomer |
|---|---|---|---|
| | Run-1 | 55.7 | 77.6 |
| | Run-2 | 55.2 | 72.7 |
| $SAA_{1-76}$ | Run-3 | 55.7 | 79.1 |
| | Average | 55.5 (0.3) | 76 (3) |
| | Run-1 | 51.2 | 69.0 |
| | Run-2 | 50.8 | 68.7 |
| $SAA_{1-104}$ | Run-3 | 51.6 | 69.1 |
| | Average | 51.2 (0.4) | 69.2 (0.4) |

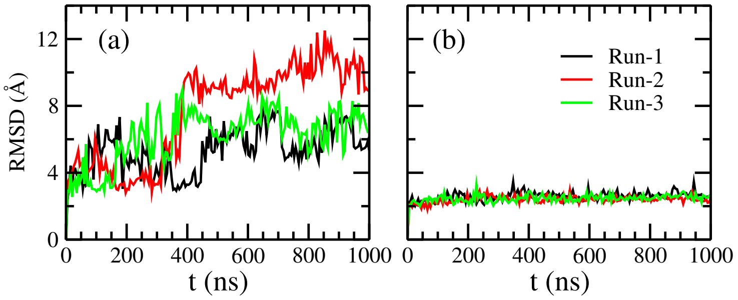


Figure SF1: Time evolution of the root mean square deviation (RMSD) of isolated (a) $SAA_{1-76}$ and (b) $SAA_{1-104}$ monomers, taking into account all non-hydrogen atoms in the first 76 residues. Data are shown for all three trajectories of each system. The RMSD values are calculated with respect to the corresponding start configurations.

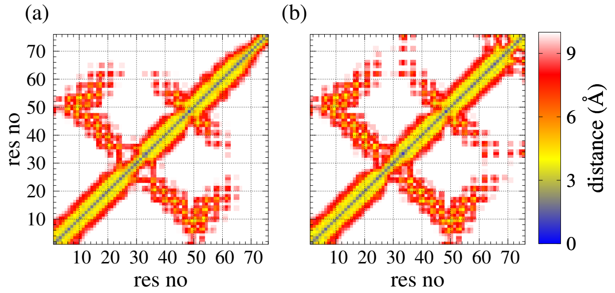


Figure SF2: Side-chain–side-chain contact map of (a) $SAA_{1-76}$ and (b) $SAA_{1-104}$ monomers as calculated over the last 500 ns of all three trajectories of each system. Data are shown only for the first 76 residues. Residue pairs whose average contact distance is more than 10 Å are excluded.

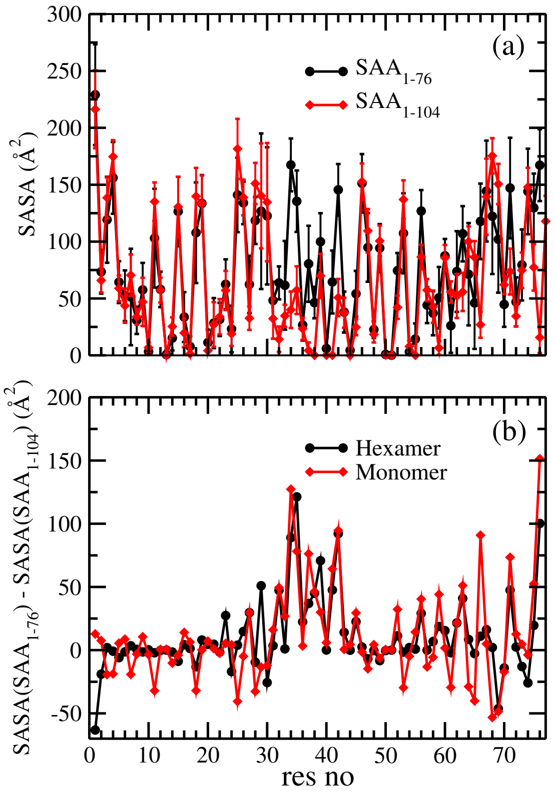


Figure SF3: (a) Average residue-wise solvent accessible surface area (SASA) of $SAA_{1-76}$ and $SAA_{1-104}$ monomers as calculated over the last 500 ns of each of the three trajectories of each system. Data are shown only for the first 76 residues. In (b) we show for each residue the difference $\Delta SASA = SASA(SAA_{1-76}) - SASA(SAA_{1-104})$ calculated either for the isolated monomer or measured in the corresponding hexamers.
